# Inadequate dried blood spot samples for Early Infant Diagnosis, how common and what are the reasons for rejection in Zimbabwe?

**DOI:** 10.1101/502500

**Authors:** Charles Chiku, Maria Zolfo, Mbazi Senkoro, Mzwandile Mabhala, Hannock Tweya, Patience Musasa, Fungai D. Shukusho, Exervia Mazarura, Angela Mushavi, Douglas Mangwanya

## Abstract

**Background:** Early infant diagnosis (EID) of HIV in infants provides an opportunity for early detection of the infection and early access to Antiretroviral treatment (ART). Dried Blood Spot (DBS) samples are used for EID of HIV-exposed infants, born from HIV positive mothers. However, DBS rejection rates have been exceeding in Zimbabwe the target of less than 2% per month set by the National Microbiology Reference Laboratory (NMRL). The aim of this study was to determine the DBS samples rejection rate, the reasons for rejection and the possible associations between rejection and level of health facility where the sample was collected.

**Methods:** Analytic cross-sectional study using routine DBS samples data from the NMRL in Harare, Zimbabwe, between January and December 2017.

**Results:** A total of 34.950 DBS samples were received at the NMRL. Of these, 1291(4%) were rejected and reasons for rejections were: insufficient specimen volume (72%), missing request form (11%), missing sample (6%), cross contamination (6%), mismatch information (4%) and clotted sample (1%). Samples collected from clinics/rural health facilities had five times likelihood to be rejected compared to those from a central hospital.

**Conclusion:** Rejection rates were above the set target of 2%. The reasons for rejection were ‘pre-analytical’ errors including labeling errors, sample damage, missing or inconsistent data, and insufficient volume. Samples collected at primary healthcare facilities had higher rejection rates.

## Introduction

Prevention of mother-to-child transmission (PMTCT) of HIV is one of the most an important tool for global elimination of pediatric HIV infection [1, 2]. WHO recommends Early Infant Diagnosis (EID) to be performed as part of PMTCT on HIV-exposed infants within four to six weeks of age [1].

In February 2013, Zimbabwe’s Ministry of Health and Child Care (MoHCC) mandated the implementation of lifelong ART for all pregnant and breastfeeding HIV positive mothers regardless of CD4 count, called Option B+[3]. This policy change represented a paradigm shift in the implementation of PMTCT and ART programs. However, only half (51%) of the HIV-exposed infants receive an EID test by the age of six to eight weeks or at earliest possible opportunity [1]. If the EID test is negative at 6–8 weeks and HIV exposure through breast feeding continues, the test must be repeated at weaning. Thereafter, definitive diagnosis after 18 months is done using rapid test[6]. It has been said that if HIV-positive infant is given ART within the first 12 weeks of life, they are 75% less likely to die from an AIDS related illness[4,5].

Dried Blood Spot (DBS) samples are preferred to the whole blood samples for EID testing as they make infant HIV testing possible even in areas with no infrastructure for collection, storage and transportation of blood samples. DBS samples are collected by pricking the heel of infants using blood lancets, drip it onto five DBS card’s spots (see fig 1), place it on dry and dust-free surface for two to four hours to allow it to dry, before packaging and send it to the National Microbiology Reference Laboratory (NMRL) through courier service.

NMRL is one of the few laboratories in Zimbabwe that tests early EID DBS samples using Roche AmpliPrep/CobasTaqMan 96 analyzer with technological capability of analyzing one full spot protocol for testing. Box 1 outlines NMRL DBS rejection criteria.

### Box 1: Zimbabwean National Microbiology Reference Laboratory (NMRL) rejection criteria

1. Incomplete identification on the requisition and/or DBS card
2. DBS card without request form/Request form without DBS card
3. Specimens with evidence of contamination, leakage or spillage in transit
4. DBS sample containing blood clots or clumps
5. DBS sample without at least 3 full spots (insufficient)

The laboratory rejects all samples that do not meet the criteria indicated in Box 1.

Figure 1 below shows the DBS samples that were being accepted for testing, anything that was not meeting this criteria in terms of volume was being rejected.

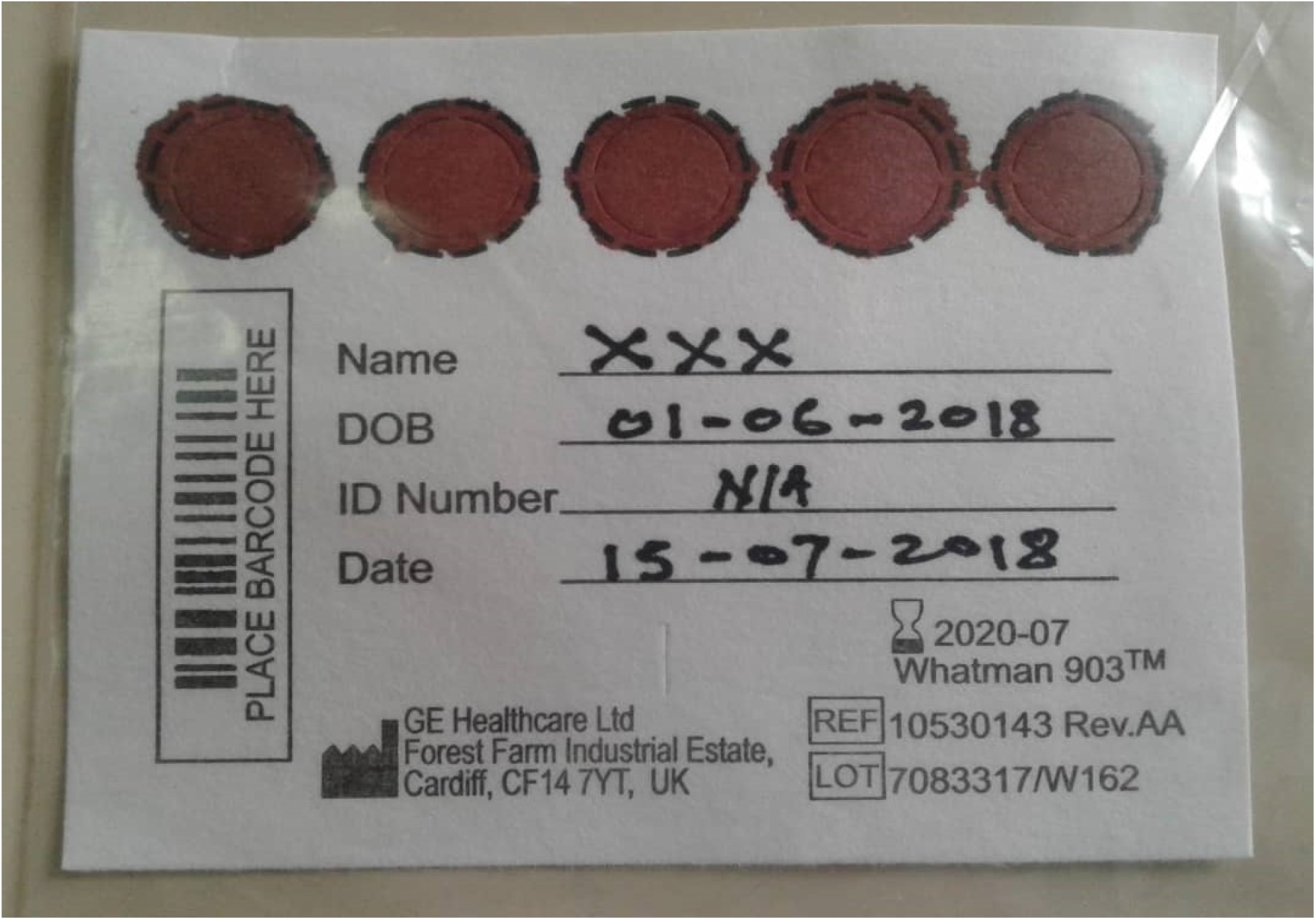
Figure 1: Sufficient dried blood spot.

There is evidence that insufficient specimen volume is one of the major reasons for high proportion of DBS rejections. It is unclear why that is a case, given that the evidence also suggests that the technology used for testing DBS samples can analyse and produce conclusive results from samples with minimum of two full spots [6]. The implication of this is that EID-eligible infants may miss diagnostic opportunities (MDO) due to DBS sample rejections. The insufficient sample volume is part of wider issues surrounding DBS sample rejection including –poor sample collection techniques at health facilities, poor documentation of samples and request forms, and samples’ failure to yield conclusive results at the laboratory.

These issues are not unique to the Zimbabwean PMTCT and ART programs; comparable programmes in other countries in Africa reported same problems. They estimated that sample collection errors contribute 60–70% towards sample rejection [7]. To take few examples, the results of a study carried out in South Africa showed that 3.7% of samples were rejected due to ‘pre-analytical’ errors including labeling errors, sample damage, missing or inconsistent data, and insufficient volume [8]. Similarly, in a study conducted in Nigeria, the main reasons for rejection were: poor collection technique (26.3%), improper labeling (16.4%) and insufficient blood collection (14.8%) [9]. The same stuy also showed that DBS collected at primary and secondary health care facilities were two to three times more likely to be rejected than those collected at tertiary healthcare facilities [10].

In Zimbabwe, it takes up to fourteen days from sample collection before NMRL communicates rejection to the submitting healthcare facility [11]. It takes further few weeks before the healthcare facility informs the mother-baby pair that the sample has been rejected and that therefore there is a need to provide a new DBS sample [10]. This inevitably results in delay in diagnosing and initiating treatment for the infants confirmed to be HIV-infected and result in MDO and loss to follow up.

There have been concerns that there is no laboratory surveillance in Zimbabwe to monitor and track EID related processes from pre-analytical phase of laboratory processes. In addition, there are very few studies that investigate the causes of DBS rejections[6]. The absence of the data on DBS rejection rates means that there is a limited intelligence to act and take appropriate corrective actions to improve the EID program and reduce loss to follow up.

The aim of this study was to determine the proportion of DBS samples that were sent to the Zimbabwean NMRL and rejected between January 2017 and December 2017 and the reasons for their rejection.

## METHODS

### Study Design

A cross- sectional study that used routine data from EID laboratory information management system (LIMS).DBS samples collected from all the facilities from five provinces of Zimbabwe referred to NMRL were logged into EID laboratory information system (LIMS). Records on rejected samples in the EID LIMS were validated against source hard copy documents. Descriptive analysis was performed to describe the variables in relation to number and proportion of rejected DBS, reasons for rejection and levels of health facility.

### Study Setting

#### General setting

Zimbabwe is a country in Southern Africa, with a population of 17 million people in 2017 [11]. The country is divided into 8 provinces and 2 metropolitan provinces (Bulawayo and Harare).

It has four tiered system of health care services, which includes (i) the primary health care facilities (predominantly rural health centers and poly clinics), (ii) district health facilities (which also includes mission hospitals), (iii) provincial hospitals and (iv) tertiary referral or central hospitals.EID samples from health facilities in five provinces (Harare, Mashonaland West, Mashonaland East, Mashonaland Central and Midlands Provinces) are tested at NMRL. The results are thereafter returned to the facility for care and treatment of the infants.

#### Specific Setting

The NMRL is an accredited laboratory situated in Harare, Zimbabwe. It was established in 2007 as the first in Zimbabwe to perform HIV DNA PCR. Until 2013, NMRL was the only laboratory service responsible for processing the DBS samples for EID, nationally. In 2013, the Zimbabwean government decentralized laboratory services and two more laboratories, Mpilo and Mutare, which now process EID samples from health facilities closer to them. NMRL only processes EID samples from the five provinces. This study analyzed DBS sample rejection rate of the samples sent to the NMRL from these five provinces. Box 1 shows the criteria for rejection.

DBS samples that met the above criteria were rejected and the laboratory sent a communication to the health facility detailing that the DBS was rejected and reason for rejection stated, in turn the health facility communicates with the caregiver of the infant to come back for another sample collection.

### Study population and Period

DBS samples collected from HIV-exposed infants from all the facilities in the five provinces referring samples to NMRL from January 2017 to December 2017 were included in the study.

### Data Collection and Validation

Data on rejected samples from the EID laboratory information management system was validated against source documents of the rejected DBS samples that were kept as hard copies. The extraction of data was done cumulatively and disaggregated by health facility on a monthly basis. Variables collected included, date of sample receipt; laboratory request number of DBS sample; total number of DBS samples received; total number of samples that were rejected; number of samples that were rejected by health facility level; health facility name, level, district, and province of rejected DBS samples; DBS rejection reason.

## DATA ANALYSIS AND STATISTICS

Descriptive analysis was conducted to describe the variables in relation to number and proportion of rejected DBS, reasons for rejection and the level of the referring health facility. The chi square test was performed using STATA version 13 (Stata Corp, Texas USA)and presented as odd ratios (OR) with 95% confidence intervals (CI). Differences at 5% level were regarded as significant

## ETHICAL CONSIDERATION

<u>Ethics approval</u>: Permission to conduct the study was obtained from National Microbiology Reference Laboratory Director, the Union Ethics Review Committee (reference number EAG/07/18) and the Medical Research Council of Zimbabwe (Ethics approval reference number MRCZ/E/194).

## RESULTS

Between January and December 2017, 34.950 DBS samples were received at the NMRL, 1291 (4%) samples were rejected. Table 1 shows the proportion of DBS samples rejected by month. The proportions of rejected samples ranged from 3% to 6% with the highest rejection rate observed in Septembe**r**.

**Table 1:** Number and proportions of rejected DBS samples at the National Microbiology Reference Laboratory, Zimbabwe from January 2017 to December 2017.

|  |  |  | 193 |
| --- | --- | --- | --- |
| **Dried Blood Spot samples** |  |  |  |
|  |  |  | 194 |
|  | **Received** | **Rejected (n, %)** | 195 |
| Total | 34950 | 1291 (4) | 196 |
| Months |  |  |  |
| January | 4138 | 105 (3) | 197 |
| February | 2959 | 98 (3) |  |
| March | 4123 | 128 (3) | 198 |
| April | 3253 | 114 (4) | 199 |
| May | 3533 | 132 (4) |  |
| June | 3150 | 107 (3) | 200 |
| July | 2668 | 106 (4) |  |
| August | 2593 | 105 (4) | 201 |
| September | 2191 | 125 (6) | 202 |
| October | 2377 | 125 (5) |  |
| November | 2189 | 88 (4) | 203 |
| December | 1776 | 58 (3) | 204 |

Table 2 shows proportion of DBS samples rejected and the association by level of health facility. DBS samples collected from clinic or rural health centers, district/faith based hospitals and provincial hospital were more likely to be rejected compared those from central hospital.

**Table 2:**
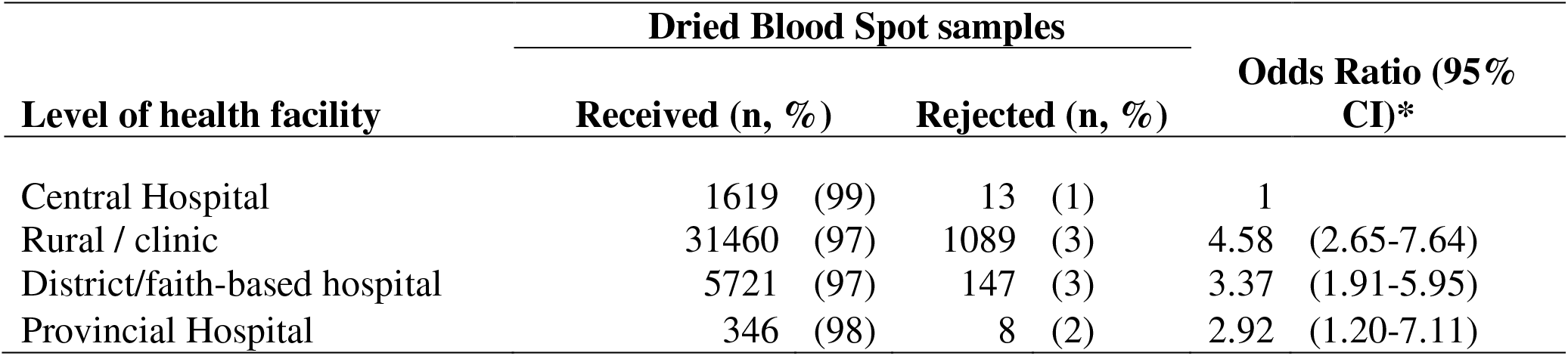
The associations between rejection of DBS samples and level of health facility from which they were collected and sent to the National Microbiology Reference Laboratory, Zimbabwe from January 2017 to December 2017.

| Level of health facility | Dried Blood Spot samples |  | Odds Ratio (95% CI)* |
| --- | --- | --- | --- |
|  | Received (n, %) | Rejected (n, %) |  |
| Central Hospital | 1619 (99) | 13 (1) | 1 |
| Rural / clinic | 31460 (97) | 1089 (3) | 4.58 (2.65-7.64) |
| District/faith-based hospital | 5721 (97) | 147 (3) | 3.37 (1.91-5.95) |
| Provincial Hospital | 346 (98) | 8 (2) | 2.92 (1.20-7.11) |

Thirty-four DBS samples that had no stated rejection reason mentioned on the form were excluded from this analysis. Majority of the DBS samples were rejected due to insufficient specimen volume. If blood did not fill all five spots on the DBS card it was considered as an insufficient volume and was rejected (Table3). Other reasons for rejections were: missing request form, missing sample, cross contamination, clotted sample, or mismatch information.

**Table 3:**
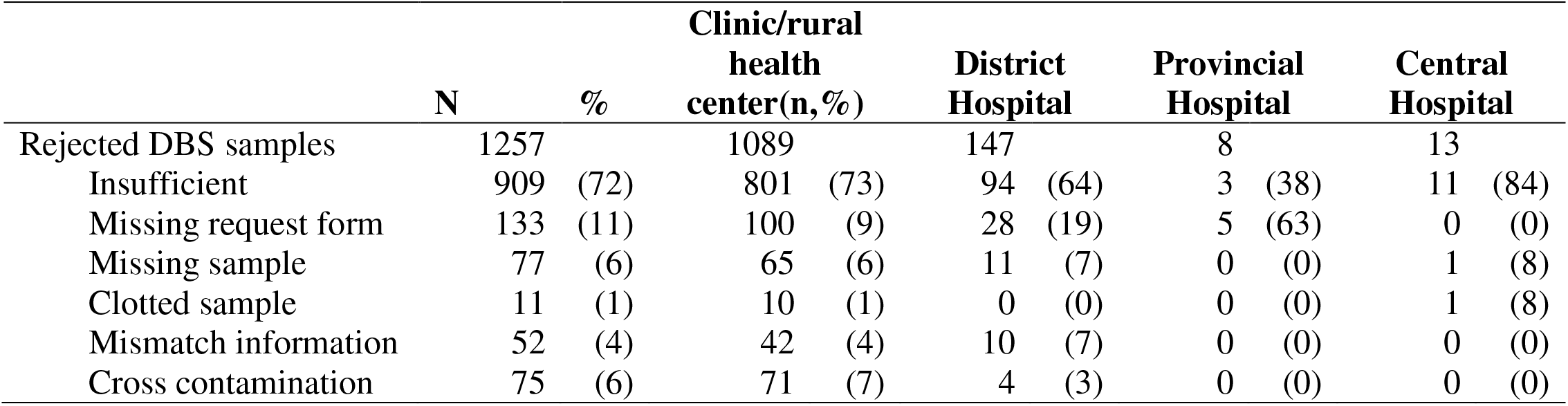
Proportion of dried blood spot samples rejected: rejection reason and type of health facility at the National Microbiology Reference Laboratory, Zimbabwe, January 2017 to December 2017.

Figure 1 shows the numbers and proportions of rejected DBS samples by province. Rejection rate ranged from 13% to 28% among the five provinces. Provinces with the highest rejection rates were Mashonaland West (28%) and Mashonaland East Province (26%).

**Figure 1:**
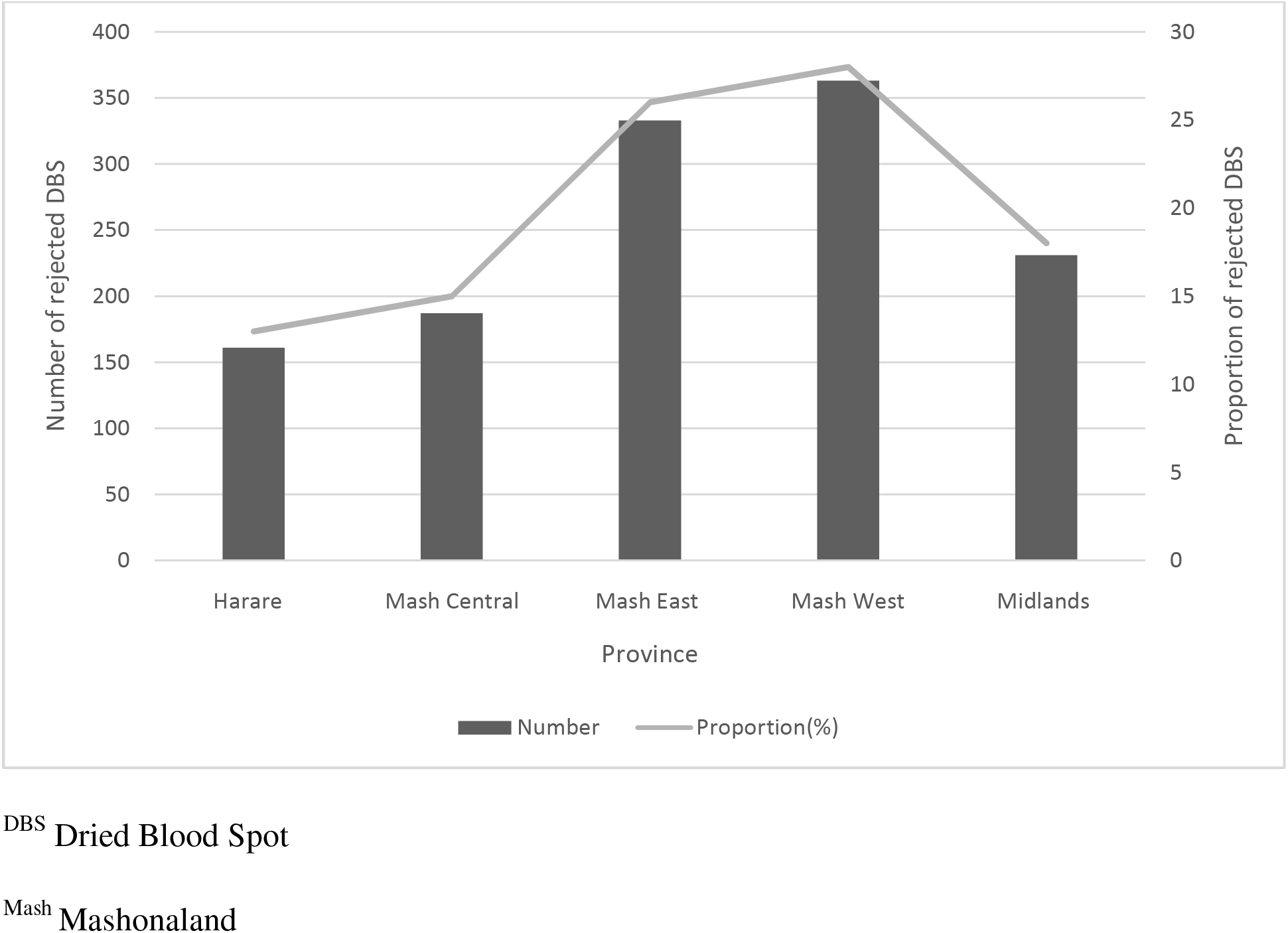
Proportion of rejected DBS samples and Provinces at National Microbiology Reference Laboratory, Zimbabwe, January to December 2017.

## Discussion

This is the first study in Zimbabwe’s EID program to assess the magnitude and reasons for rejection of DBS samples at NMRL, which analyzes samples referred by five provinces. The national maximum rejection target for DBS samples is<2% per month. This study found that the NMRL’s rejection rate is above the national average. These findings were similar to the findings of another study conducted in Kwazulu Natal province in South Africa where the rejection rate was 4% [6]. Even higher aggregated EID program DBS rejection rate of 7.4% was reported in similar study conducted in Zimbabwe in 2017 in Mashonaland West province [10]. However, the latter was a smaller study looking at only one province compared to the current study that looked at five provinces.

Analysis showed that samples that were collected at primary healthcare facilities (rural clinics) level facilities where most of the patients received services were five times more likely to be rejected. Findings were similar to a Nigerian where most rejections were from facilities that serves most patients [9]. One possible explanation for this higher rejection rate in primary healthcare facilities is the high demand and the low financial resources for staff recruitment and equipment available in these sites, which might result in reduction of quality assurance mechanisms on sample collection.

The other possible explanation could be that those rural facilities are unable to offer adequate in-service training to members of staff. They can afford to small number staff for training in large urban centers with the view to cascade training the rest of the teams. The research by Smit *et al* reported that in-service skills cascade is not effective as model of training. Those receiving onsite training often receive poorer quality of training than their counterparts do. Smit *et al* goes on to posit that staff orientation and mentorship on DBS collection, storage and transportation is essential in standardizing the skills and improving DBS sample collection [12]. Their conclusions are consistent with findings from Nkengasong’s, study which showed that sensitizing health care workers on activities of sample collection, handling and completion of laboratory request forms resulted in reduction in errors from 19.05% to 6.76%[13].

The current study found that large proportion of rejection were due to insufficient blood volume collection. These findings were consistent with the results of the study in South Africa, where 48% of all rejection was due to insufficient sample collection [6]. A lower proportion 14.8% of rejection due to insufficient blood collection was observed in another Nigerian study [7].

A previous study has shown that the use of two insufficient spots of DBS samples for the analysis yields satisfactory results. Govender *et al* showed that the use two insufficient spots, prevented 10.504 samples from being rejected due to insufficient blood volume. However, there was no denominator available in this study, for us to determine the proportion of prevented rejections [6].

However Govender*et al* conducted a validation study on the modified method of testing to determine if the use of two insufficient DBS spots protocol can yield the same results as a validated method of one full spot protocol [6]. This study showed that the two insufficient spot protocol yielded results that are comparable to the validated one full spot protocol. These findings create a need for NMRL to revise its rejection criteria in relation to the emerging evidence and consider accepting samples with two full spots.

Furthermore, there has been evidence that implementation of quality management system improves identification of shortfalls of the system and corrective actions taken eliminates the root causes of the problems, thereby reducing the rejection rate of samples [14, 15].

This study has found that Mashonaland West Province contributed the highest proportion of rejected DBS samples, followed by Mashonaland East Province. Further, investigation is required to establish the possible causes.

The implications of the rejections may result in missed diagnostic opportunities of HIV-infected infants, loss to follow up on the infants and challenges in early initiation of HIV-positive infants on ART. DBS rejection rates contribute towards delays in accessing of laboratory results for EID testing, have serious implications on the PMTCT program and lives of infants that may be in need of life long ART[8,10].

## Limitations

The EID program does not have a unique identifier for patients, therefore, we could not track if another DBS sample was collected. This resulted in challenges to determine the extent of delay in the collection of another sample.

We could not observe the staffing levels and workload of the health facilities that were sending samples to NMRL. Additionally, we could not determine the qualifications, competencies and trainings on DBS collection of the personnel that were collecting DBS samples, which might have an impact on quality assurance mechanisms for the sample collection

There was no available data to track how long it takes from sample collection to communicating the test results to the mother-infant pair. This could have shown whether rejection has an effect on infants getting early start of ART.

## Recommendations

- NMRL to monitor the rejection rates and notify the supervisors of health facilities in real time so that corrective and preventive actions are taken to avoid delays due to rejected DBS.
- Revision of user handbook to make it clear that blood drop must spread along the DBS card till it reaches the marked sections of the circle.
- Develop mentorship programs on DBS sample collection, storage and transportation especially in areas where there are higher rejection rates.
- District laboratories or facilities with laboratories could evaluate the samples for quality before sending to NMRL, since they may pass through district lab first. This would allow deficiencies to be identified closer to the health facility from where samples are collected, and actions could be taken at district level before sending the samples to the NMRL.
- Laboratory processes needs to strengthen implementation of quality management system to identify deficiencies in DBS sample management so that corrective actions are taken to continuously reduce sample rejection rates that impacts patient care.
- NMRL needs to conduct a laboratory validation of “two insufficient spot protocol” against a” one full spot protocol” so that the rejection criteria may be reviewed based on the scientific evidence of the validation.

## Conclusion

This study has shown that DBS rejection rates were above the national target. The major reason for rejection was insufficient volume samples. Clinic/rural health centers have higher rejection rate than central hospitals. Over all, there is need to monitor rejection rates in real time, so that corrective and preventive actions may be taken to reduce or eliminate causes of DBS rejection.

## Acknowledgements

This research was conducted through the Structured Operational Research and Training Initiative (SORT IT), a global partnership led by the Special Programme for Research and Training in Tropical Diseases at the World Health Organization (WHO/TDR). The training model is based on a course developed jointly by the International Union Against Tuberculosis and Lung Disease (The Union) and Medécins sans Frontières (MSF). The specific SORT IT program which resulted in this publication was implemented by: Medécins Sans Frontières, Brussels Operational Centre, Luxembourg and the Centre for Operational Research, The Union, Paris, France. Mentorship and the coordination/facilitation of these SORT IT workshops were provided through the Centre for Operational Research, The Union, Paris, France; the Operational Research Unit (LuxOR); AMPATH, Eldoret, Kenya; The Institute of Tropical Medicine, Antwerp, Belgium; The Centre for International Health, University of Bergen, Norway; University of Washington, USA; The Luxembourg Institute of Health, Luxembourg; The Institute of Medicine, University of Chester, UK; The National Institute for Medical Research, Muhimbili Medical Research Centre, Dar es Salaam, Tanzania.

## Funding

The programme was funded by: the United Kingdom’s Department for International Development (DFID); La Fondation Veuve Emile Metz-Tesch supported open access publications costs. The funders had no role in study design, data collection and analysis, decision to publish, or preparation of the manuscript.

## Conflict of interest

None declared.

